# Grapevine scion gene expression is driven by rootstock and environment interaction

**DOI:** 10.1101/2023.01.12.523795

**Authors:** Zachary N Harris, Julia E Pratt, Laszlo G Kovacs, Laura L Klein, Misha T. Kwasniewski, Jason P Londo, Angela Wu, Allison J Miller

## Abstract

**BACKGROUND:** Grafting is a horticultural practice used widely across woody perennial crop species to fuse together the root and shoot system of two distinct genotypes, the rootstock and the scion, combining beneficial traits from both. In grapevine, grafting is used in nearly 80% of all commercial vines to optimize fruit quality, regulate vine vigor, and enhance biotic and abiotic stress-tolerance. Rootstocks have been shown to modulate elemental composition, metabolomic profiles, and the shape of leaves in the scion, among other traits. However, it is currently unclear how rootstock genotypes influence shoot system gene expression as previous work has reported complex and often contradictory findings.

**RESULTS:** In the present study, we examine the influence of grafting on scion gene expression in leaves and reproductive tissues of grapevines growing under field conditions for three years. We show that the influence from the rootstock genotype is highly tissue and time dependent, manifesting only in leaves, primarily during a single year of our three-year study. Further, the degree of rootstock influence on scion gene expression is driven by interactions with the local environment.

**CONCLUSIONS:** Our results demonstrate that the role of rootstock genotype in modulating scion gene expression is not a consistent, unchanging effect, but rather an effect that varies over time in relation to local environmental conditions.

## Background

Grafting is an ancient horticultural technique that joins genetically distinct organ systems to generate chimeric individuals [1–3]. Most frequently, grafting is used to fuse together the root system of one individual, which becomes the rootstock, to the shoot system of a different individual, the scion. Grafting has been used in at least 70 major woody perennial crops to confer favorable traits to trees and woody vines such as dwarfing, changes in the timing of fruit ripening, increased fruit yield and quality, and resistance to biotic and abiotic stress [2]. Ongoing research aims to understand how grafting and different rootstock genotypes impact function and phenotype of the scion.

Among the most notable applications of grafting was its use to save the European grapevine (*Vitis vinifera* ssp. *vinifera*) from the North American aphid-like insect, phylloxera, that was introduced to Europe in the mid-1800s [4]. Native North American grapevine species (*Vitis* spp.) have co-evolved with phylloxera and can tolerate infestation by impeding insect damage in most of their root system. European grapevine varieties, on the other hand, have no such natural tolerance and are susceptible to the insect feeding off its roots, leaving wounds that ultimately cause death of the vines. Today, phylloxera is nearly ubiquitous, and the cultivation of *V. vinifera* in areas where phylloxera exists is possible only as a scion grafted on a phylloxera tolerant rootstock. In addition to tolerance to phylloxera, grafting has also been used to adapt elite European grapevine scion cultivars to various environmental conditions [5–7], and today, at least 80% of vineyards worldwide comprise European grapevine scions grafted to North American *Vitis* species [8]. Despite the myriad ways in which grafting has been used to aid grapevine cultivation, the extent to which rootstock genotype modulates scion phenotype remains a topic of intense investigation. Recent studies have shown that rootstock genotypes influence shoot elemental composition, leaf shape and vigor [9–13] and that there are subtle influences of rootstock on the metabolome of leaves in the grafted scion [12, 14, 15]. However, key questions remain in terms of how rootstock genotype influences scion gene expression.

Several studies have sought to understand the role of grafting on grapevine scion gene expression, but results are complex and sometimes contradictory. One general question is whether the physical act of grafting induces changes in gene expression, and if so, whether those changes reflect the genotype of the rootstock. In a comparison between Cabernet Sauvignon grafted to a different species (a heterograft) and self-grafted controls (homografts), the graft junction of heterografts showed differential expression of genes related to stress response and plant defense within a month after grafting [16]. Four months after grafting, shoot apical meristems of grafted Cabernet Sauvignon showed differential regulation in genes that impact chromatin modification and hormone signaling, among other functional categories [17]. No differentially expressed genes were identified across comparisons of different rootstock genotypes for heterografted individuals, suggesting that the observed changes in gene expression were a result of heterografting and not from specific genome-genome interactions. This result was further supported by studies of Chambourcin scions in which vines grafted to different rootstocks exhibited few differentially expressed genes as a function of rootstock genotypes [10, 12]. In contrast, a study of grafted Gaglioppo reported that >17,000 genes were differentially expressed in leaves of scions grafted to different rootstock genotypes [18]. This suggests that under certain conditions, rootstock genotypes elicit distinct transcriptomic differences in heterografted vines.

A second set of questions on the nature of rootstocks modulating the scion transcriptome addresses whether the effects of grafting or rootstock genotype change over time (over the course of the season or across years). This question is of particular importance because these temporal factors, when included in experimental designs, tend to be the largest descriptors of variation in gene expression within a single tissue [12, 19, 20]. In a study examining how the effect of grafting changes over a season, Cabernet Sauvignon berries showed differential expression of genes related to auxin across rootstock genotypes, but the general effect was diminished as the season progressed [19]. Similarly, berries from Pinot Noir differentially expressed genes related to cell wall metabolism, stress responses, and secondary metabolism across a rootstock and irrigation experiment, but the results were diminished later in the season [21]. These results seem to indicate that differential patterns of gene expression observed in scions grafted to different rootstocks are apparent early in the season, but diminish later in the season. However, this effect was not universal. Subsequent studies in Pinot Noir showed that the rootstock effect on differential expression was stronger in mature berries than in developing berries, with particular differences noted in genes related to secondary metabolism [20]. Variation in these studies ranging from general patterns over time to scion- and rootstock-genotype-specific effects suggest that there are additional factors that may influence how rootstock genotypes shape gene expression in the shoot system.

One key factor which often confounds comparisons across gene expression studies in grapevine is variation in environmental conditions where the vines were grown. Plants exhibit transcriptomic responses to natural and seasonal environmental variation [22–24], and growth under field conditions tends to present plants with a complex combination of stress conditions [25]. However, most gene expression studies in grapevines have examined the effect of applied stress under controlled conditions rather than natural environments experienced in the field. For example, transcriptomic responses have been shown in water stress [26], salt stress [27, 28], and differential exposure to light [29]. It is not uncommon for different *Vitis* species, or even different genotypes, to display distinct transcriptomic responses to stress. For example, a cultivar of *V. amurensis* was shown to have a stronger transcriptomic response to cold stress than a cultivar of *V. vinifera* which resulted in a muted physiological response [30]. Differential gene expression in stress response has also been observed in genotypes used as rootstocks where, for example, root and leaf gene expression differentially varied across an irrigation treatment in the rootstocks M4 and 101.14 [31]. These results suggest that in grafted vines with two distinct genotypes, transcriptomic responses may vary in the rootstock genotype relative to the scion genotype. Further, transcriptomic response in one graft partner may impact how the other partner responds to a particular environmental stress or condition. As a result, grafting likely adds an additional dimension of variation in how grapevines modulate their phenotypic response to diverse environmental conditions. Namely, grafted vines have revealed ways in which the below-ground and above-ground portions of the plant respond to controlled stress conditions, and how they interact with each other, to produce dynamic phenotypic changes over time. How grafting mediates the transcriptomic response to natural environmental variation as it changes over time in the field remains an open question.

In this study, we assessed the influence of grafting, rootstock genotype, time of season, year, and local environmental conditions and their interactions on gene expression in the grapevine cultivar Chambourcin. To do this, we sampled leaf and reproductive tissues (flowers and fruits) from ungrafted (own-rooted) Chambourcin vines as well as vines where Chambourcin was grafted to one of three different rootstocks. Samples were collected at three phenological stages (anthesis, veraison, harvest-ripe) in each of three years. Through this design, we sought to answer the following questions: 1) How do grafting and rootstock genotype influence shoot system gene expression? 2) Does the influence of grafting and rootstock genotype on shoot system gene expression vary over time? and 3) Is there an environmental component to rootstock influence on shoot system gene expression? Data presented here demonstrate that the influence of rootstock genotype on shoot system gene expression is highly dependent on tissue type and time of sampling (both year and time of season), suggesting that the impact of grafting on gene expression in the scion varies over time. Follow up analyses indicate that these differences are not strictly temporally correlated, but related to the local environmental conditions that the vines are experiencing.

## Results

### Experimental Design

This study took place in a rootstock experimental vineyard located at the University of Missouri Southwest Research Station near Mount Vernon, Missouri (see [10] for a detailed description). We collected samples from 72 individuals of the grapevine cultivar Chambourcin growing ungrafted (own-rooted) and grafted to three different root systems: 1103P, 3309C, and SO4 (N = 18 vines per root/shoot combination; Supplemental Figure 1). Leaf and reproductive tissue were collected from each vine at three phenological stages (~50% anthesis, ~50% veraison, and immediately prior to the harvest-ripe stage) over three consecutive years (2017, 2018, 2019). After accounting for sample loss and low-quality extractions, we sequenced the transcriptomes of 1,178 samples.

### Sequencing counts and high-level descriptions of variation

We obtained 4.04M reads per sample (SD=1.36M) on average using the 3’-RNAseq protocol. We mapped reads to the 12Xv2 reference grapevine genome and observed 3.44M uniquely mapping reads per sample (85.08%, SD=1.15M). On average, 3.28M reads per sample aligned uniquely to gene features (SD=1.10M). Some reads were discarded due to multimapping (mean=398K, SD=149K) or because they did not align to gene features (mean=148K, SD=80K). Gene counts were normalized using DESeq2 and filtered such that only genes with counts greater than four in at least four samples were retained, resulting in a data set with 24,392 genes measured in 1,178 samples. Principal component analysis (PCA) on 24,392 genes showed that the first two PCs captured 19.5% and 12.4% of the total variation, respectively (Figure 1). In PC space, leaf samples clustered together and reproductive samples formed two distinct clusters (Figure 1A). Within the tissue clusters, there was clear structure from year (Figure 1B) and phenological stage (Figure 1C). There was no clear rootstock signal on the first two PCs (Figure 1D).

**Figure 1:**
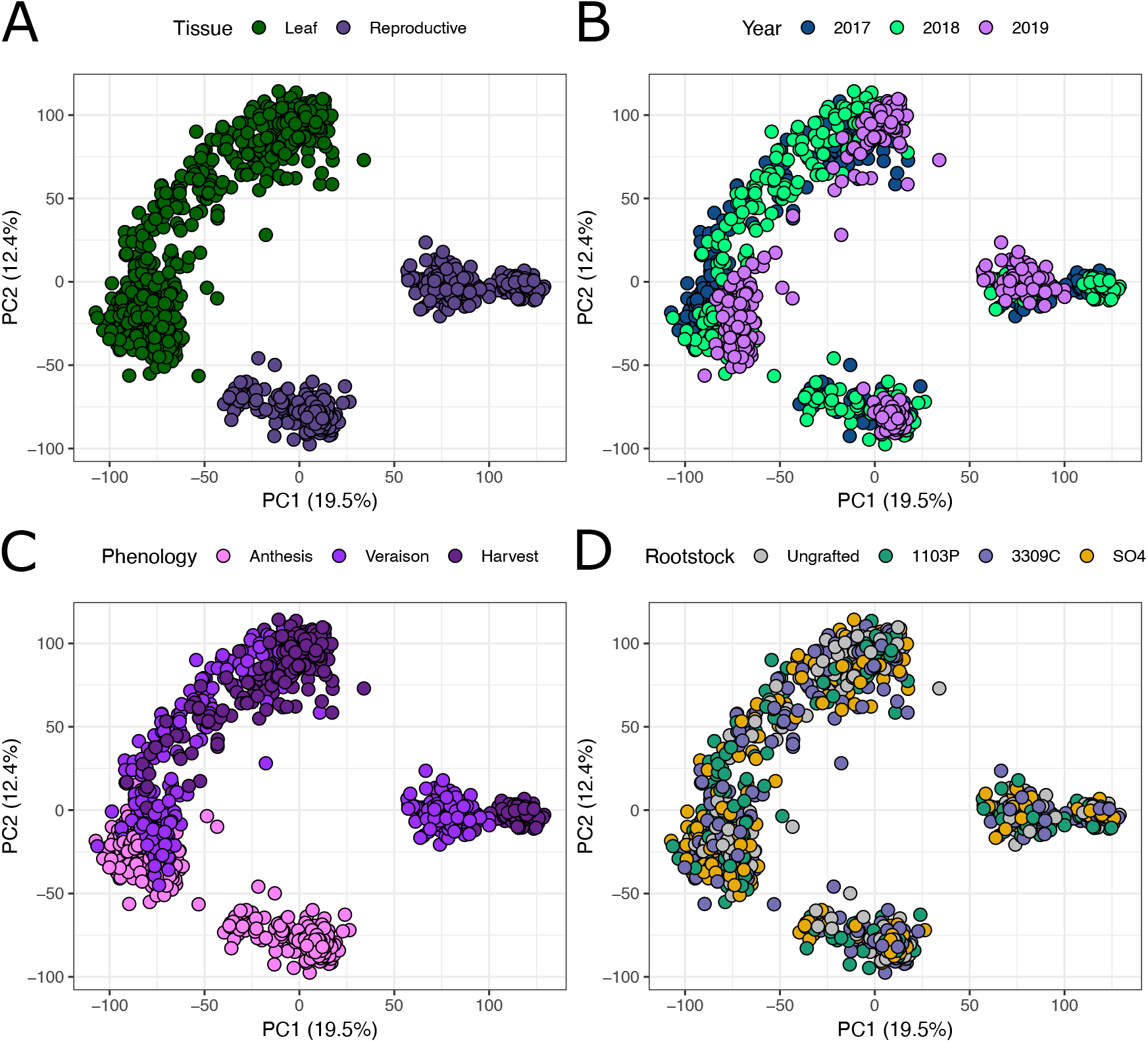
PCA on gene expression colored by tissue, year, phenology, and rootstock The top two principal components of the quality filtered, normalized, and VST-transformed gene counts, as colored by **A)** tissue, **B)** year of sampling, **C)** phenological stage, and **D)** rootstock genotype.

### Self-Organizing Maps for rootstock main effect

From the PCA and our previous work in our rootstock experimental vineyard [10, 12], we predicted that the rootstock main effect would be subtle. In order to investigate rootstock effects on scion gene expression, we fit linear models to each measured gene after transforming each gene’s expression with a variance stabilizing transformation. The expression of each gene was modeled with rootstock genotype, tissue, year, and phenological stage as main effects and with all pairwise interactions. Irrigation was included in the model, but was not formally interpreted as it was previously found to be of negligible effect [12, 32]. Each linear model was evaluated under a variance explained framework, and genes in or above the 75th percentile (0.44%) of variance explained by rootstock were retained. The resulting set of 5,495 genes was used to train a self-organizing map (SOM) to identify genes responding similarly in Chambourcin tissues across rootstocks (Figure 2). The SOM was trained to identify 81 clusters (9 by 9 hexagonal grid), of which 51 had at least 16 genes and were significant for the rootstock main effect in post-clustering linear models. For comparison purposes, the relationship between the SOM and PCA are provided (Figure 2A-B).

**Figure 2:**
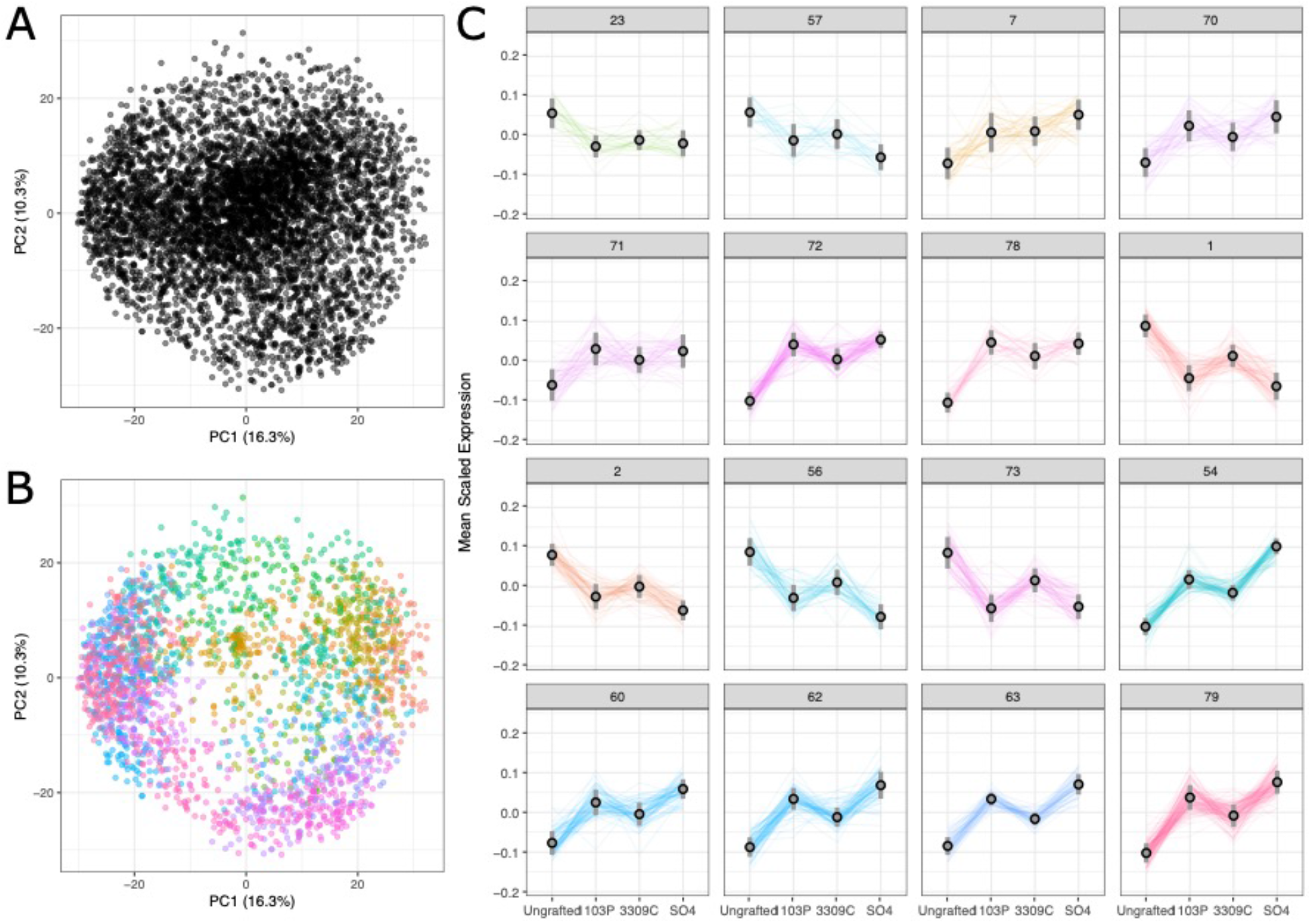
Self-organizing map captures clusters of genes that vary with rootstock genotype across three years of study **A)** A principal components analysis on all genes across the samples showing low-dimensional embeddings of variation in scion gene expression. **B)** The principal component plot, colored by assignment to SOM clusters and filtered for proximity to the median gene in the cluster to show the relationship between SOM and PCA. **C)** Examples SOM clusters that showcase commonly occuring patterns. Mean scaled expression for genes assigned to example SOM clusters (numbered) that were significant for rootstock in post-clustering linear modeling are shown.

From all clusters identified by the SOM, several key patterns in gene expression point to consistent effects of grafting, as well as rootstock specific effects (Figure 2C). For example, we identified sets of genes that were consistently down regulated in grafted vines relative to ungrafted vines (clusters 23, 57) and a separate set of genes that were consistently upregulated in grafted vines relative to ungrafted vines (clusters 7, 70, 71, 78). None of these clusters were significantly enriched for any functional categories. In addition, we observed rootstock genotype-specific effects on gene expression patterns in the scion. The most prominent patterns were clusters in which expression was more similar in leaves of Chambourcin grafted to 1103P and SO4 than it was to ungrafted vines or 3309C-grafted vines. Within the clusters representing this most common pattern, expression was sometimes higher in ungrafted vines (clusters 1, 2, 56, 73). Cluster 73 was enriched for a single functional category (‘cysteine-type endopeptidase inhibitor activity’, GO:0004869). In other clusters, expression was lower in ungrafted vines (clusters 54, 60, 62, 63, 79). Cluster 54 was enriched for the functional categories ‘cytosolic ribosome’, (GO:0022626), ‘structural molecule activity’ (GO:0005198), and ‘cellular amide metabolic process’ (GO:0043603). Cluster 62 was enriched for the functional categories ‘ribosome’ (GO:0005840), ‘ribonucleoprotein complex’ (GO:1990904), and ‘structural constituent of ribosome’ (GO:0003735). Cluster 63 was enriched for the functional categories ‘structural constituent of ribosome’, ‘structural molecule activity’, and ‘ribosome’.

### The influence of rootstock genotype in a tissue-specific, time-informed analysis

The SOM identified a clear but subtle signal of rootstock genotype on the scion transcriptome. To further understand what in our experiment explains this observation and why it has been missed in previous studies, we performed a traditional analysis of differential expression using DESeq2. Traditional analyses with DESeq2 allowed us to analyze each rootstock comparison across tissues and across each of the time points (phenology and year) within our study. In general, few genes were identified as differentially regulated across rootstock genotype in leaves or reproductive tissue at any time point in 2017 or 2019 (Figure 3A). However, in 2018, comparisons across rootstock genotypes showed many differentially expressed genes across all three phenological stages. The largest number of differentially expressed genes were identified in comparisons between ungrafted and grafted vines; rootstock genotype specific patterns of gene expression were less prominent. This pattern is especially apparent at later phenological stages (Figure 3A). In general, more genes were up-regulated in grafted Chambourcin than were down-regulated. Overall, the differences due to rootstock were very subtle (Figure 3B). For example, if genes were filtered to only consider comparisons with a log2 fold change larger than two, the number of genes dropped by 54% to 93% for pairwise comparisons between ungrafted and 1103P-grafted vines.

**Figure 3:**
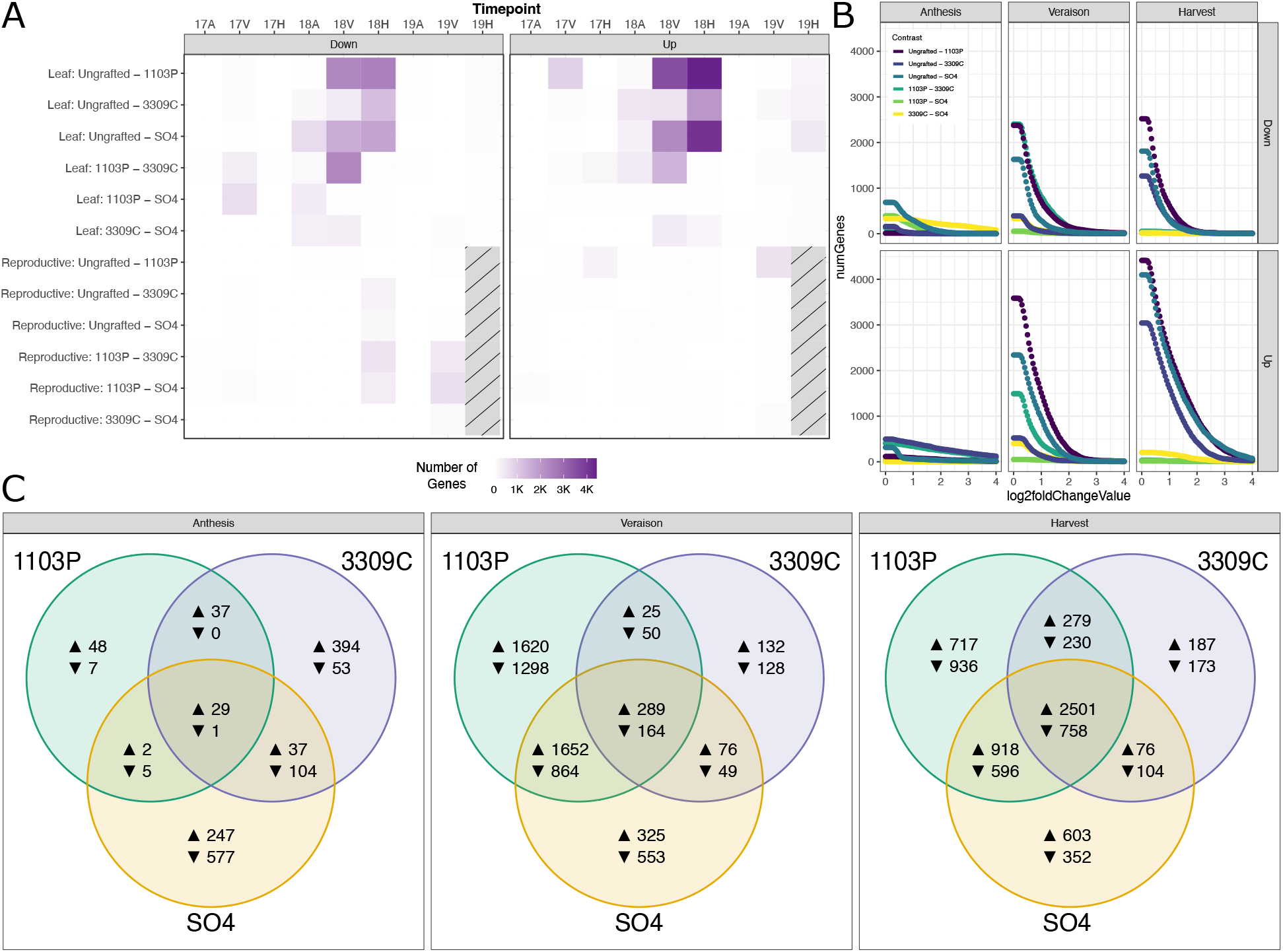
Differentially expressed gene counts are enriched for a single year of study **A)** A heat map showing the number of genes identified as differentially expressed across rootstock contrasts, broken down by tissue, year, phenology, and direction of change (17A = 2017 anthesis, 17V = 2017 veraison, etc). Genes characterized as differentially regulated are presented in reference to the rootstock on the right (in the comparison labeled “Ungrafted - 1103P”, genes designated as ‘Up’ are more highly expressed in 1103P). **B)** Effects size scans showing the number of genes we would retain (y-axis) if we were to filter on various log2 fold-change thresholds (x-axis) within 2018 leaves. **C)** Venn diagrams comparing grafted vines to ungrafted vines in 2018 leaves across phenological stages. Genes upregulated in grafted vines are shown next to an up arrow, where genes down-regulated in grafted vines are shown next to a down arrow.

Leaves from Chambourcin vines grafted to 1103P and SO4 were more likely to have unique functional categories of genes enriched when compared to ungrafted vines (Supplemental Table 1). For example, in 1103P-grafted vines, there were 129 genes differentially regulated at anthesis, 5,962 genes differentially regulated at veraison, and 6,935 genes differentially regulated at harvest relative to ungrafted vines (Figure 3B-C). Functional categories uniquely enriched in the comparison between ungrafted and 1103P-grafted vines were only identified at veraison where a suite of functions related to general cellular growth and activity were upregulated, including “cellular macromolecule biosynthetic process” (GO:0034645), “peptide biosynthetic process” (GO:0043043) and “amide biosynthetic process” (GO:0043604). Similarly, leaves from SO4-grafted vines showed 1,002, 3,972, and 5,908 differentially regulated genes at anthesis, veraison, and harvest, respectively, relative to ungrafted vines. Several functional categories were enriched in anthesis in SO4-grafted vines including those related to protein formation, such as “peptide biosynthetic process” (GO:0043043), “translation” (GO:0006412), and “amide biosynthetic process” (GO:0043604). Interestingly, we note a strong suite of functions down-regulated in SO4-grafted vines at veraison related to ungrafted vines including “gene expression” (GO:0010467), “nucleic acid metabolic process” (GO:0090304), “nucleobase-containing compound metabolic process” (GO:0006139), “RNA metabolic process” (GO:0016070), and “RNA processing” (GO:0006396). Vines grafted to 3309C generally had fewer unique differences in gene expression when compared to ungrafted vines. However, several functions were enriched among down-regulated genes in 3309C at anthesis, mostly related to telomere maintenance and DNA conformational changes, including “telomere maintenance via telomerase” (GO:0000722), “telomere capping” (GO:0016233) “DNA geometric change” (GO:0032392), and “DNA duplex unwinding” (GO:0032508).

While the individual rootstock genotypes elicited some unique responses in the scion transcriptome (Figure 2C), many genes were influenced by multiple rootstocks when compared to ungrafted vines. For example, at veraison, one of the largest effects on the transcriptome came from the overlap between ungrafted and 1103P-grafted vines and ungrafted and SO4-grafted vines where 1,652 genes were jointly upregulated, and 864 genes were jointly downregulated in the grafted vines (Figure 3C). Functional analysis of the upregulated genes showed enrichment for terms related to ‘microtubule-based process’ (GO:0007017), ‘microtubule-based movement’ (GO:0007018), and ‘movement of cell or subcellular component’ (GO:0006928). At harvest, we observed a large number of genes differentially regulated across all three rootstock genotypes relative to ungrafted. Here, we identified 2,501 shared genes that were up-regulated relative to ungrafted and 758 genes that were down-regulated relative to ungrafted. Only the up-regulated gene set contained enriched functionality, many of which were shared in veraison, including ‘microtubule-based process’, ‘microtubule-based movement’, ‘movement of cell or subcellular component’, and ‘cytoskeleton organization’ (GO:0007010).

### Environmental Analyses

The unique signature of rootstock genotype on scion gene expression identified in 2018 prompted us to consider what in 2018 differed from the rest of our study. An on-site weather station captured 10 features of the local environment, reporting hourly measurements of average temperature, total precipitation, wind speed, average relative humidity, average solar radiance, total radiation density, pressure, average dew point, estimated reference crop evapotranspiration, and calculated clear sky solar radiation. These hourly measurements were used to build 26 composite statistics representing the minimum value, maximum value, change in value, and mean value for most features over a 24-hour window. Precipitation and radiation density were summed (rather than averaged) to build a composite statistic. Composite statistics were built for every day for the three years of this study to test the correlations across features. Given that many environmental features were highly correlated, we opted to collapse this variation using a principal components analysis (hereafter called the environmental or ePCA), from which we extracted data for each of the nine days of sampling for subsequent analyses. The top two ePCs explained a total of 61.9% of the environmental variation. The first ePC (41.3%) primarily captured variation in mean values of temperature, pressure, and solar radiation (Figure 4A). The second ePC (20.6%) captured variation in temperature, humidity, and radiance stability and variation in mean pressure and humidity.

**Figure 4:**
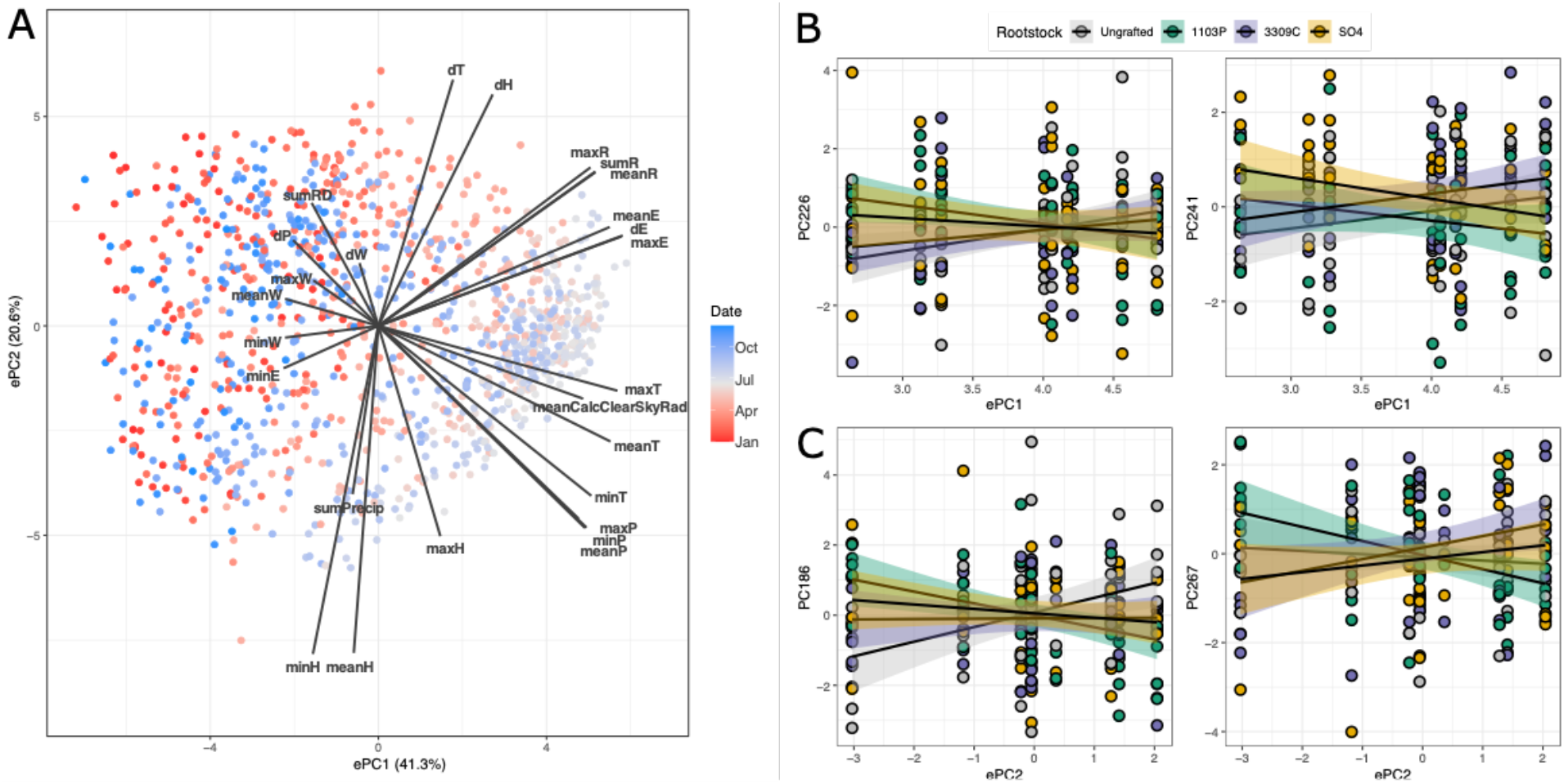
The environmental PCA and its relationship to gene expression as mediated by rootstock genotype **A)** A PCA biplot showing the span of environmental variation over the course of three years and how the features of the environment load onto those PCs. **B)** Gene expression PCs (gPCs) significant for the interaction of rootstock and the first environmental principal component, ePC1. **C)** gPCs significant for the interactions of rootstock and the second environmental principal component, ePC2.

In order to understand the influence of the environment on gene expression, we summarized variation in gene expression using PCA (hereafter called the gene expression PCA or gPCA). In the gPCA, 288 gPCs explained 80% of total variation in the transcriptome. Each gPC was fit with a linear model parameterized with each ePC as a main effect and in interactions with tissue and rootstock. For each gPC, the environment was considered significant if at least 5% of the variation was explained by the environment or an interaction with the environment. Briefly, ePC1 explained significant variation in 10 gPCs as a main effect and 11 gPCs through the interaction with tissue. In each case, the interaction between the environment and tissue were characterized by crossing slopes (as opposed to slopes that were just different in the same direction) indicating that leaves and reproductive tissue were responding to the environment in different ways (Supplemental Figure 2). When considering ePC2, nine gPCs were significant for the environment main effect, and nine gPCs were significant for the environment by tissue effect.

In addition to responding to the environment main effect and the tissue by environment interaction, some gPCs varied significantly with the rootstock by environment interaction. For example, gPC226 and gPC241 were significant for the interaction of rootstock and ePC1 (Figure 4B). In both cases, vines that were ungrafted and vines that were grafted to 3309C had positive associations with ePC1 while 1103P- and SO4-grafted vines had negative associations. Similarly, gPC186 and gPC267 were significantly associated with ePC2 as modulated by rootstock, but these patterns of association were quite variable (Figure 4C). For example, gPC186 was positively associated with ePC2 in ungrafted vines, while all grafted vines had negative associations with ePC2. Similar but distinct patterns were reflected in correlations between gPCs and ePCs 3-4. In total, 12 gPCs were influenced by the interaction of rootstock genotype and the environment. We looked to see if genes that loaded heavily (>1.96 sd away from the average loading) on the gPCs were significantly enriched for functional roles. Of the 12 gPCs influenced by interaction of rootstock genotype and the environment, six had exactly one term enriched in either highly loading genes or lowly loading genes: “RNA modification” (GO:0009451). To gain a higher resolution insight to the broad classification, we looked to see if any protein domains (Pfam and InterPro) were similarly enriched in the gene sets. Only considering the top two ePCs, many of the domains enriched on the gPC loadings were similar. For example, the Pfam domain ‘NB-ARC’ (PF00931) was enriched on genes loading positively on PC226 (significant for ePC1) and PC186 (significant for ePC2). gPC226 had five Pfam domains that were enriched in negatively loading genes, including “reverse transcriptase-like”, “reverse transcriptase”, “retrotransposon gag protein”, or “RNase H-like domain found in reverse transcriptase”, and a domain of unknown function, “transposase-like DUF 659” (PF13456, PF00078, PF03732, PF17919, and PF04937, respectively). Similar domains were also enriched in gPC241 (significant for ePC1) and gPC267 (significant for ePC2). Other gPCs significant for the interaction of rootstock and the environment were additionally enriched for domains associated with the “PPR repeat family” (PF13041) and “DYW domain” (PF07727).

## Discussion

In this study, we showed that rootstock genotype influences gene expression in the scion of grafted grapevines. This influence was demonstrated as a general effect and through interactions with tissue, time, and the local environment. This work supports previous results suggesting that rootstock genotypes have a measurable effect on the scion phenotype in grafted plants. Our results indicate that rootstock effects, even when subtle, are complex, manifesting in particular tissues at particular time points, likely through interaction with the local environment.

### Rootstock influences scion gene expression independent of tissue, phenology, or year

Previous work in grapevine has demonstrated that grafting and rootstock genotypes can alter gene expression of the scion, but the complete profile of this effect has been difficult to piece together. For example, heterografting alters gene expression of Cabernet Sauvignon tissue agnostic to rootstock genotype [16, 17], while leaves of the cultivar Gaglioppo showed substantial variation in a rootstock-genotype-specific manner [18]. Moreover, previous work in Chambourcin at a single time point showed virtually no differential expression by rootstock genotype or graft status, suggesting these effects are not ubiquitous [10]. Trends over time are even less clear [12, 19, 20]. Potential reasons for these discrepancies include the use of different scion-genotype pairs, difficulty in identifying subtle differences across rootstock genotypes, and disparate environmental conditions. In the present study, we focused on the latter two potential causes.

Given the prior expectation of very small effect sizes, we employed self-organizing maps (SOMs) to identify clusters of genes that respond similarly across samples and can then be understood both functionally and in the context of the experimental design (Figure 2). We showed that many genes were subtly responding to rootstock genotype, and that their responses can be grouped into various patterns. The most common pattern was that gene expression of Chambourcin leaves grafted to 1103P and SO4 were often quite similar to each other and distinct from ungrafted and 3309C-grafted vines. While these efforts have increased our capacity to interpret functional differences between effects of rootstock genotypes on scion gene expression, few functional categories were identified in the clusters we observed. This could be explained in two ways. First, the responses we identified were due to a general effect that did not have any particular functional role. This is possible as both 1103P and SO4 are considered to be vigor-inducing rootstocks, meaning that they tend to allocate more resources to scion foliar growth than to scion reproductive effort [33]. In contrast, 3309C is considered to be a low-vigor rootstock, with less dramatic foliar resource allocation. Clusters of genes with strong expression influence from the rootstock could just be highlighting these differences by genome-wide differential gene regulation. Second, more advanced techniques to represent meaningful embeddings of high-dimensional data are still in their infancy and are especially underexplored in the context of plant gene expression data. For example, there is currently no commonly employed method to learn an optimal grid size for SOMs, which would allow for the generation of more refined clusters that could have functionally identifiable roles. Techniques like variational autoencoders could aid in refining the functional understanding of this effect, but the software that could perform this task efficiently is only recently being developed [34, 35]. Regardless, the persistent identification of patterns seen previously in other phenotypes [32], despite their functional interpretation, suggested that our samples contained a signal that was not previously observed in the Chambourcin transcriptome. This warranted deeper analysis in the context of temporal and environmental variation.

### Rootstock differentially influences gene expression over time

Substantial effort has been devoted to characterize growth and development of grape berries [36]. Collectively, this work showed that there is a clear developmental program in grapevine, but that there can be variation in that program [37–39]. Recent studies have examined the impact of grafting on berry development, but reported conflicting results across systems and environments [19, 20]. Here, analyzing rootstock contrasts in different tissues across three phenological stages for three years revealed a strong temporal effect on rootstock modulation of the leaf transcriptome. In particular, we only see notable differentially expressed genes in leaves sampled in 2018 (Figure 3). We note this effect becomes stronger as the season progresses, supporting the results of Zombardo et al. [20]. During anthesis in 2018, we observe only a handful of genes differentially expressed by rootstock (as compared to ungrafted), with larger numbers observed for genes down-regulated in SO4-grafted vines (577) and genes up-regulated in 3309C-grafted vines (394). At veraison, many more genes were differentially expressed in 1103P-grafted vines and a large suite of genes was shared between 1103P- and SO4-grafted vines. By harvest, there was still considerable overlap between 1103P- and SO4-grafted vines, but the largest effect was shared between upregulated genes across all grafted genotypes relative to ungrafted vines. Functionally, this gene set was enriched for intracellular movement, including microtubule-based processes, cytoskeleton organization, and cell cycle processes. These results suggest that while there were differentially expressed genes in the grafted scion at particular times in the season due to unique to rootstock genotypes, the largest effect was likely a general response to grafting at the end of the season. The fact that this result was only observed in 2018 suggests that the vines were experiencing different conditions in that particular year which elicited a rootstock-mediated response. A similar effect was observed in a series of vineyards in Europe where the year of sampling was the largest descriptor of variation in the transcriptome [40]. An identical effect was previously observed in shoot elemental composition [41] and could indicate an interaction between rootstock and the vine’s local environment.

### Scion gene expression varies across the rootstock by local environmental interaction

Plants growing under field conditions experience a range of environmental conditions that trigger stress responses throughout the growing season [25, 42]. Responses to environmental variation can be detected in multiple phenotypes [41, 43–45], are often highly complex, and in general are not predictable from laboratory experiments [25]. In fact, variation in the local environment can influence the expression of well-studied molecular pathways, such as flowering [46], disease resistance [47], and circadian rhythm [48]. A recent study on gene expression in maize inbred lines showed that even variation in microclimates across a single field led to variation in expression of 15% of the maize transcriptome[45]. Understanding this variation is vital to deciphering the basis of physiological changes across a season and to predict the impacts of global climate change on plant growth. This is especially important in grapevines where the effects of climate change are predicted to be substantial [49, 50] and are already being observed [51]. However, to our knowledge, this is the first study investigating the role of rootstock genotype in the scion transcriptomic response to environmental variation.

In the three years of this study, we observed that gene expression in grafted Chambourcin scions varied with the local environment: many gene expression principal components (gPCs) were highly correlated with environmental principal components (ePCs). Of note, ePC1 explained significant variation in 10 gPCs, and ePC2 explained significant variation in nine gPCs. Multiple gPCs were additionally influenced by rootstock × environment interaction; for example, ePC1 interacted with rootstock genotype to explain variation in gPC226 and gPC241. In both cases, the slopes of associations between environment and transcriptome were more similar in 1103P- and SO4-grafted vines (both negative slopes) as compared to ungrafted and 3309C-grafted vines (both positive slopes). As noted above, this pattern was also frequently observed in associations between shoot element composition and the environment in the same vineyard [41]. Across all gPCs that have significant variation explained by the interaction of rootstock and the environment, we identified only one gene ontology term enriched on genes loading strongly to the gPCs: “RNA modification” (GO:0009451). RNA modification is a GO term with many child terms including RNA base conversion (substitution), RNA base insertion, and RNA base deletion. However, we only observed the broad category to be enriched. In order to understand this effect, we carried out enrichment analyses for other functional information including Pfam domains and Interpro accessions. Functional domains most likely to be enriched in this analysis included the NB-ARC domain, domains related to retrotranscription and retrotransposition, and PPR and/or DYW domains.

In plants, NB-ARC domains are associated with R genes, common in pathogen defense response [52]. At a minimum, this suggested that scions grafted to different rootstocks exhibit different defense responses in the scion, which has been reported in grapevine and many other woody perennials [1, 2, 4]. However, this also suggests that environmental variation present at a single site exerts differential pathogen pressure on the vines over time. This is unsurprising as the conditions necessary for some grapevine pathogens can vary over time in a single vineyard [53]. Genes related to retrotranscription also being enriched in this analysis could lend support to this hypothesis. Retrotranscription is a common function of retroviruses during infection, and the differential regulation of NB-ARC domain-containing genes could be responding to such infections. However, given the simultaneous enrichment of terms related to retrotransposition, it is more likely that variation in the environment is driving changes in the activation of retrotransposons. Transposons are known to be environmentally responsive and have a predisposition to target genes related to environmental response [54]. However, how this effect is modulated by rootstock genotype requires further work. Finally, genes with DYW and PRR domains typically associate with RNA editing in organellar transcripts, most commonly through C to U conversions [55]. This is an important avenue for future experiments given that organellar transcripts tend to dominate the cellular mRNA landscape [56]. More work is needed to understand the functional implications of these genes being influenced by the interaction of rootstock and the environment.

### What underlies phenotypic variation in perennial clonally propagated, grafted plants?

Our previous work identified phenotypes in the scion of grafted grapevines that vary significantly with rootstock genotype, including leaf elemental composition [32], leaf shape [10, 12], and berry chemistry [57]. Where the transcriptome can be thought of as a coordinated system to maintain optimal performance in real time, these other phenotypes may reflect cumulative, season-long, perturbations to vine activity. In short, these phenotypes may reflect a record of the vine’s past experience. That we can identify rootstock effects in these phenotypes, but see little difference in real time transcriptomic responses, may indicate that the genomic underpinnings of these responses are not manifest at the transcriptional level, but at a higher order level. Given that we have a clonally replicated scion, differences due to genomic sequence variation in the scion are unlikely. However, data presented here provide some evidence to suggest that previously observed phenotypic differences may be due to variation in the epigenome of the scion. First, the genes differentially expressed in this study point to variation in the activity of transposons and RNA base conversion, both of which are epigenomic processes. Moreover, we show here that early in the season, 3309C elicited down-regulation of genes related to DNA geometric conformation and telomere maintenance, also connected to epigenetic processes. Finally, a recent study showed that vines grafted to a single rootstock, 3309C, maintained different patterns of DNA methylation than ungrafted vines [58]. Together, these studies may point to changes in the scion epigenome as one potential mechanism underpinning the ubiquitous nature of rootstock influence on shoot system phenotype.

## Conclusions

In the present study we show that the influence of rootstock genotype on scion gene expression is dynamic, displaying variation over time and in association with local environmental conditions. We observe that some clusters of genes tend to have subtle variation across all time points, but the lack of functional information likely highlights general effects from vigor induction of some rootstock genotypes. However, large effects are only observed at particular time points when local environmental conditions are atypical. We showed that in 2018, when the environmental conditions were different from the other years of this study, many differentially expressed genes could be identified. Interpreting our gene expression results in the context of this environmental variation showed several genes expressed in the scion were modulated by the interaction of rootstock genotype and the local environment. Such observations could explain why previous studies have found contradictory results: there is likely a large influence from the local environment on rootstock modulation on scion gene expression. Moving forward, studies should be designed to uncover subtle general results or to capture a large range of environmental variation to further tease apart the complex nature of rootstock influence on scion gene expression.

## Methods

### Study Design

Samples were collected from a rootstock experimental vineyard managed by the University of Missouri’s Southwest Research Center in Mount Vernon, Missouri, USA (37.074167 N; 93.879167 W) (Supplemental Figure 1). This vineyard has been used extensively to measure variation in leaf morphology [10, 12], berry and leaf metabolomics [12, 57], leaf elemental composition [10, 12, 41], and vine physiology [59] across different rootstock scion combinations. This vineyard features the hybrid grapevine cultivar Chambourcin growing ungrafted (own-rooted) and grafted to three commercially available rootstocks: 1103P, 3309C, and SO4. Each Chambourcin/rootstock combination was planted in replicated blocks of four vines per row per rootstock/scion combination for nine rows. From each replicated rootstock/scion block, we sampled the middle two vines. From each vine, we sampled two tissue types: leaf and reproductive. For leaves, the youngest, fully-opened leaves from two shoots were pooled as a single sample per vine. For reproductive tissue, we sampled either unopened flower buds (early season, anthesis) or berries (veraison and harvest), which were similarly pooled by vine. Samples were collected in row-order from 10:00AM to approximately 2:00PM. Samples were immediately flash frozen in liquid nitrogen and were transported to the lab where they were stored in a −80°C freezer. Samples were collected from three phenological stages: anthesis (~50% flower buds open), veraison (~50% of berries turned from green to red), and immediately prior to harvest. Samples were collected in three years: 2017, 2018, and 2019. Berry samples were not collected from harvest 2019 as powdery mildew rendered most fruit unharvestable.

### Extraction and Sequencing

To maximize the number of samples sequenced in this study, we opted to perform a reduced-representation approach to RNAseq called 3’-RNAseq, which performs well in organisms with reasonably characterized genomes [60]. For this procedure, total RNA was extracted from each tissue using the Spectrum Plant Total RNA kit (Sigma-Aldrich, St. Louis, MO, USA) with 2% PVP40 added to the extraction buffer to sequester phenolic inhibitors. Extractions were checked for quality using a Nanodrop (Thermo Fisher Scientific, Waltham, MA, USA) and sequenced using an NextSeq500 (Illumina, San Diego, CA, USA). The resulting data set contained single-end, 86 base pair reads.

### Differential Expression Analysis

Samples with fewer than 500,000 reads were discarded. Low-quality reads were removed based on the overrepresentation of *k*-mers using BBduk (April 11, 2019 [61]. Reads were then aligned to the 12Xv2 reference genome [62] using STAR v2.7.2b [63] with default alignment parameters. Reads aligning to annotated gene features were counted using featureCounts v2.0.1 [64] against the VCost.v3 reference grapevine genome annotation [62]. Due to potentially mis-annotated gene boundaries, the annotation was modified to extend gene regions 500 bp. Differential expression analysis was carried out in DEseq2 v1.26.0 [65]. Each gene was modeled with each of the following main effects: block, irrigation, tissue, year, phenology, and rootstock. Genes with normalized counts less than four in fewer than four samples were removed, and the gene-wise dispersions were re-estimated. This model and variance stabilizing transformed data [66] were saved for future use.

After the initial fit, experimental metadata (tissue, year, phenology, and rootstock) were concatenated into a single composite term in order to assess higher-level interactions. Each gene was re-estimated with a model containing the concatenated metadata, irrigation, and block as fixed effects, although the effects from irrigation and block were not considered in this study. Each rootstock contrast was then analyzed within each tissue × year × phenology interaction. From these models, normalized counts (using DESeq2’s implementation of the variance stabilizing transformation) were extracted for genes mapping to two broad classes of constitutively expressed house-keeping gene families: ubiquitin-domain related (IPR: IPR000626) and actin domain (IPR: IPR004000) (Supplemental Figure 3). Variation in expression of these genes was assessed across samples for generally consistent patterns, although large changes have been reported from factors such as tissue, phenology, etc [12, 67].

### Self-Organizing Maps

Due to the complex nature of the experimental design, we wanted to thoroughly explore the rootstock main effect independent from all other sources of variation. Prior to the full differential expression analysis, we used the VST-transformed expression to fit independent linear models to scaled expression for each gene. We fit these models to include the full experimental design up to and including all two-way interactions of the following terms: tissue, year, phenology, and rootstock. All genes that had more than 75th percentile for variation explained by rootstock were used to train a self-organizing map (SOM) [68]. The SOM was used to identify genes that responded similarly across rootstocks. The SOM was trained on a 9 × 9 hexagonally-connected grid and presented with the data in 500 iterations while linearly decreasing the learning rate from 0.05 to 0.01 over the training process. Each node was considered an independent cluster of genes, and only the genes that were within the 50th percentile of distance to the node center were retained [69]. Each gene in each node (subsequently called a cluster), was summarized by taking the mean across samples. Linear models with rootstock as the only fixed effect were then applied to each cluster. Clusters that were significant for rootstock (alpha = 0.05/81) were analyzed for functional enrichment.

### Environmental Data Analysis

An onsite weather station [70] captured hourly measurements of temperature, precipitation, wind speed and direction, relative humidity, solar radiation, radiation energy density, pressure, dew point, estimated short crop evapotranspiration, and clear sky radiation. From each of these, we built composite summaries of the 24 hours preceding sampling including minimum values, maximum values, and change in values over the window. Composite statistics built from 24 hours preceding sampling were highly correlated with composite statistics built from 24 hours before sunrise on the day sampling and smaller windows including four and six hours before sampling. Moreover, many traits within the 24-hour window were highly correlated, so we collapsed the correlation structure using PCA to understand variation in gene expression as a function of broad environmental variation. We explored the top four environmental PCs (ePCs) as they collectively captured 80% of environmental variation.

Similarly, we compressed variation in gene expression using PCA. We explored the top 288 gene expression PCs (gPCs) which collectively explained 80% of the gene expression variation. For each gPC and ePC combination, we fit linear models to capture the environmental main effect, the tissue main effect, the rootstock main effect, and all possible interactions of these model terms. For this portion of the study, we focused only on the environmental main effect, the rootstock by environment interaction, the tissue by rootstock interaction, and the rootstock by tissue by environment interactions. Models were assessed under an effect size framework where all terms with more than 5% of variation were subjected to post-hoc comparisons of slopes. Where the post-hoc comparisons were significant (Tukey-adjusted p-value < 0.05), we explored the genes that loaded heavily (>1.96 sd away from the mean loading) onto the gPCs using functional enrichment analysis.

### Functional Enrichment

Enriched Gene Ontology (GO) terms were identified using gProfiler2[71]. First, gene names were mapped from the VCost.v3 names to the more broadly used 12Xv2 names. Then, a query was made using the reference organism “vvinifera” within the “annotated” domain scope. Each run was internally corrected for multiple tests using the ‘fdr’ correction. Functional enrichments within PCs and SOMs-derived gene lists were considered significant using an alpha threshold of 1e-05, while rootstock contrasts and overlaps from DESeq2 were considered significant using an alpha threshold of 4.6e-04. gProfiler was used such that only terms associated with ‘biological process’ (GO:BP) were identified. Following the enrichment analysis, GO terms were clustered by semantic similarity using Revigo (similarity=0.5) [72]. We note that despite only using terms associated with the label GO:BP, Revigo occasionally merged those terms with other categories, usually ‘molecular function’ (GO:MF).

In addition to GO term enrichment, we sought to characterize more specific functional annotations and their enrichments. The entire set of predicted gene models from the VCost.v3 genome annotation were functionally annotated using InterProScan[73]. From this functional annotation, we looked for functionally enriched terms as identified by Pfam and InterProScan with E-values < 1e-10. Enriched terms were identified using the hypergeometric test implemented in the phyper function in R. For both sets of terms, significance was assessed by comparisons of p-values to an alpha threshold corrected for the number of genes considered (Bonferroni).

## Data Availability

Sequencing data are provided on the NCBI Sequence Read Archive under the following accessions: PRJNA674915 and PRJNA915033. Two samples were found to be corrupted: 471R and 624L. Both are provided in truncated forms within PRJNA915033. All code used for the analyses of these data are provided on Github: https://github.com/PGRP1546869/mt_vernon_1719_rnaseq. Additional data including experimental metadata, environmental data, and a persistent version of record of all code are provided on FigShare: https://doi.org/10.6084/m9.figshare.21861480.

## Author Contributions

This study was conceptualized by ZNH, LLK, LGK, JPL, MTK, and AJM. Sample tissues were collected by AJM, LLK, LGK, and ZNH. Data were generated by JPL. Data were curated and analyzed by ZNH, JEP, LLK, AW, and JPL. Funding was acquired by LGK, JPL, MTL, and AJM. Methodology and data visualization were completed by ZNH and JEP. ZNH and AJM wrote the original draft. All authors contributed to reviewing and editing the final manuscript. All authors approve of this submission.

## Funding and Acknowledgements

This work was supported by the National Science Foundation Plant Genome Research Program 1546869. We thank members of the Kovacs Lab at Missouri State University and the Miller Lab at the Danforth Plant Science Center and Saint Louis University for helping to collect samples, and Hannah Martens, Amy Swezc-McFadden, Katheleen Deys from the Londo Lab at Cornell for processing samples. We additionally thank members of the Miller Lab for editing and improving this manuscript.

## Ethics Approval, Consent to Participate, and Consent of Publication

Not applicable

## Supplemental Figures

**Supplemental Figure 1:**
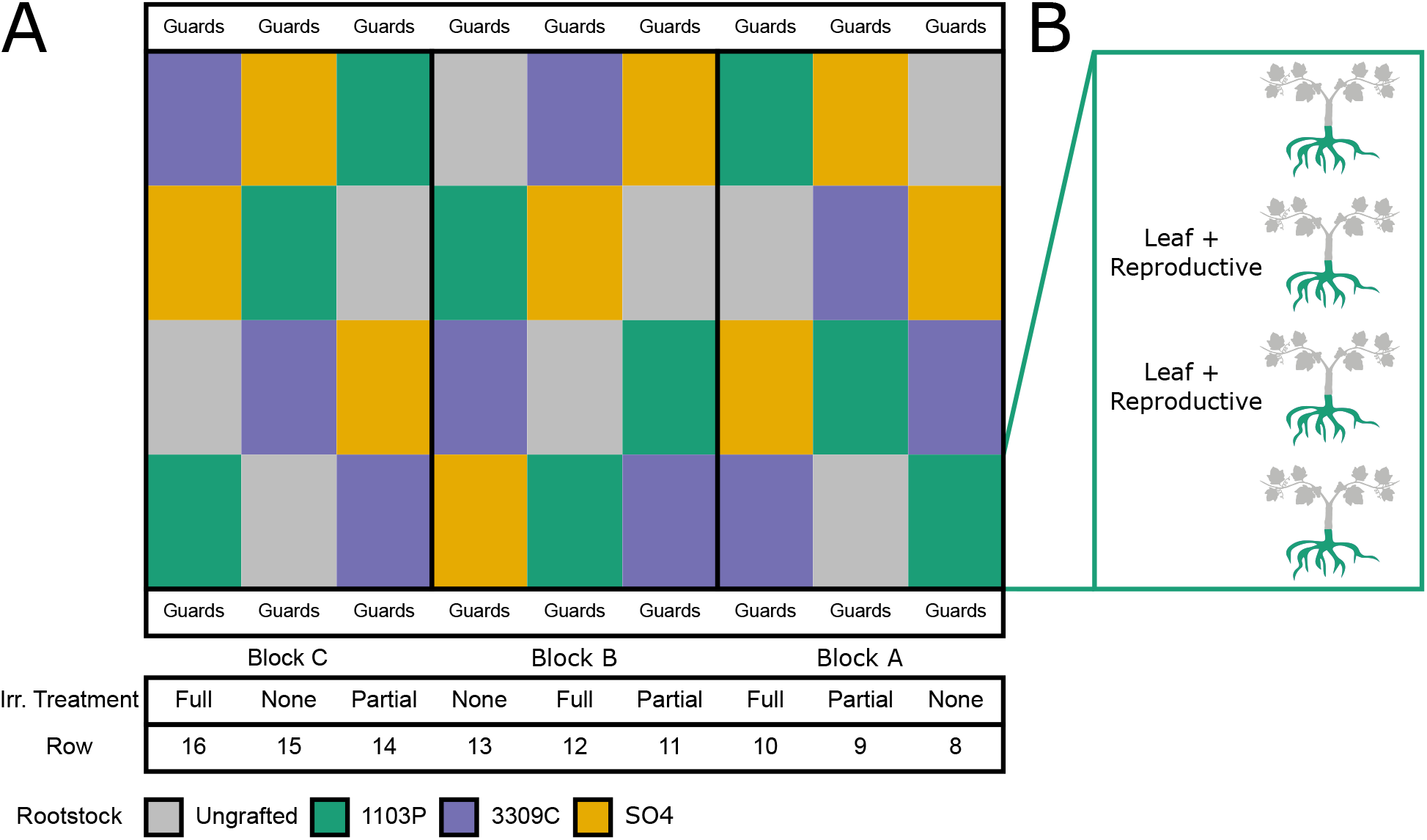
Experimental Design. **A)** Vineyard layout. The vineyard contains the grapevine cultivar Chambourcin grown ungrafted and grafted to three commercial rootstocks: 1103P, 3309C, and SO4. Each row of the vineyard contains all rootstock/scion combinations and is treated with one of three irrigation regimes: full (100% replacement of evapotranspiration), partial (50% replacement of evapotranspiration), or none (no replacement of evapotranspiration). **B)** Each cell of the vineyard features 4 replicated vines. Samples (leaf and reproductive) were collected from the middle 2 vines in each cell. This figure is partially adapted from [10], which is provided under the Creative Commons license (CC BY 4.0).

**Supplemental Figure 2:**
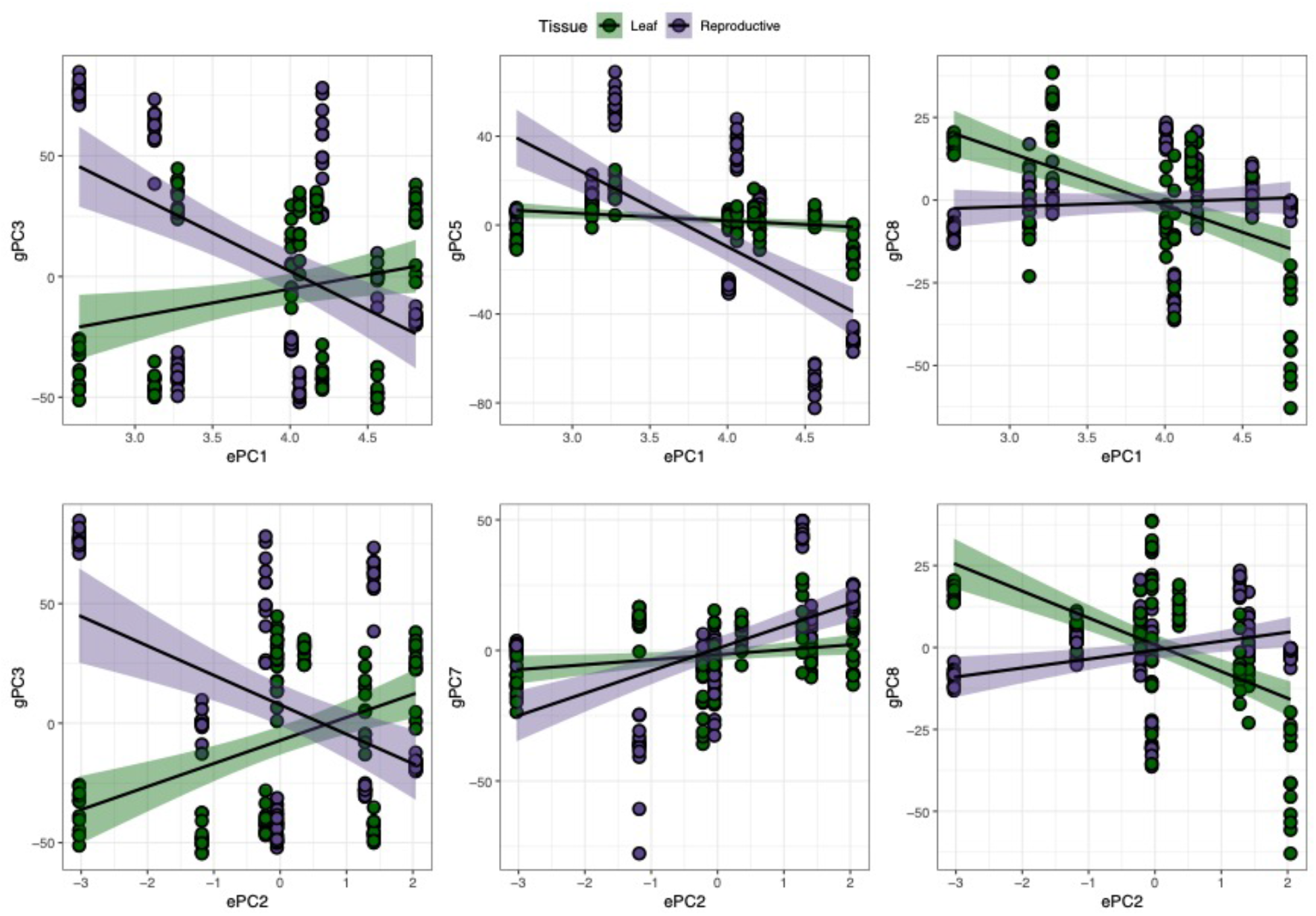
Example correlations between gene expression PCs and environmental PCs which differed by tissue. ePC1 and ePC2 are shown against the gPCs for which they explained large proportions of variation.

**Supplemental Figure 3:**
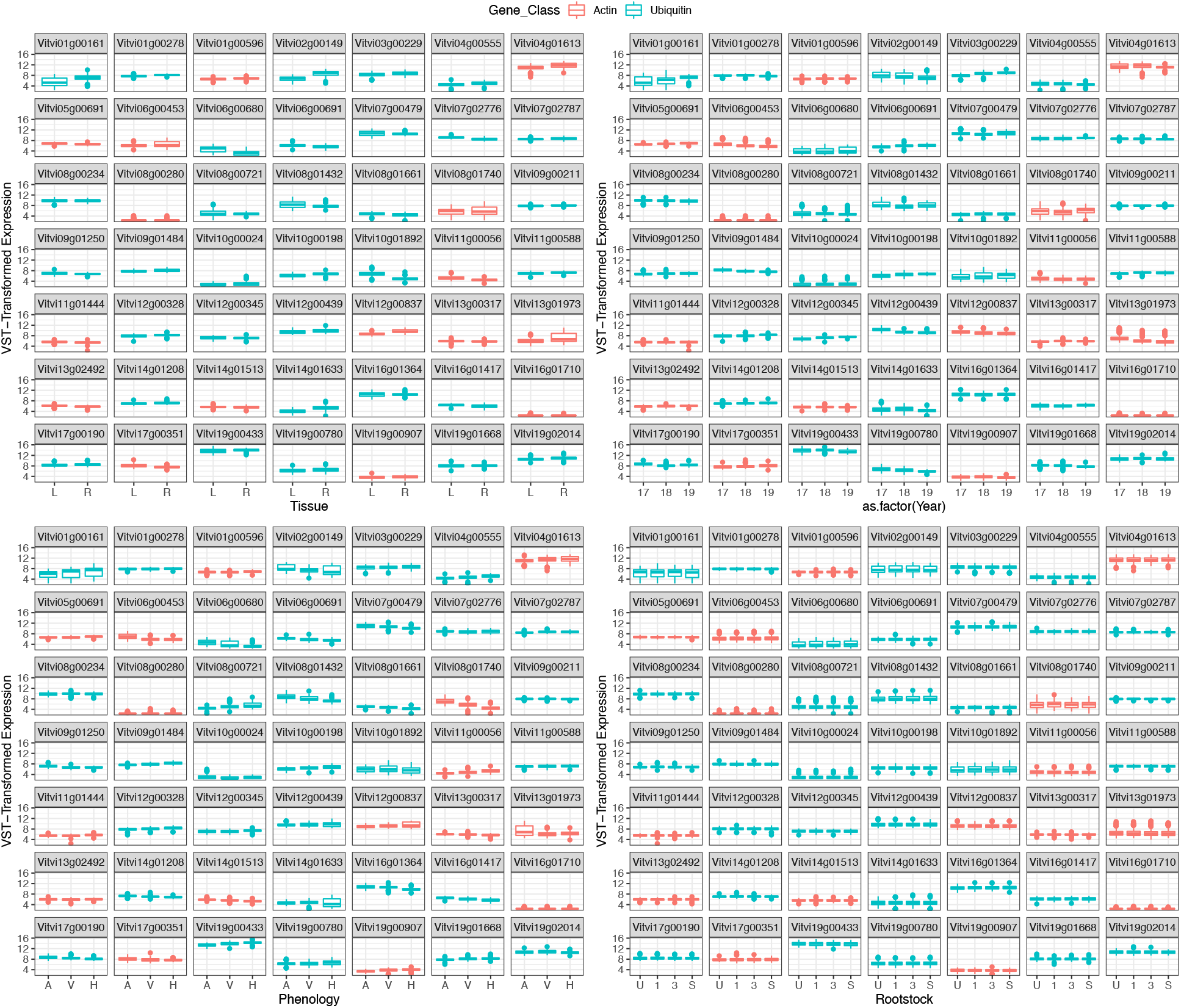
Survey of housekeeping genes. Two classes of housekeeping genes (Actin (IPR004000) and Ubiquitin (IPR000626)) were plotted against the major factors in the experiment’s design (tissue, year, phenological stage, and rootstock genotype). Factor names are abbreviated to the first character of their name (Leaf: L, Reproductive: R, Anthesis: A, Veraison: V, Harvest: H, Ungrafted: U, 1103P: 1, 3309C: 3, SO4: S).

**Supplemental Table 1:** GO terms enriched in grafted vines by rootstock as compared to ungrafted.

